# Contextual unfamiliarity during drug intake induces transient AMPA receptors linked to relapse vulnerability

**DOI:** 10.1101/2025.11.24.690243

**Authors:** Henryque Alles, Xiaojie Huang, Oliver M. Schlüter

## Abstract

Relapse to drug use remains a challenge in treating substance use disorders (SUDs). In rodent models of SUDs, Ca²⁺-permeable AMPA receptors (CP-AMPARs) in nucleus accumbens (NAc) medium spiny neurons enhance cue-drug memories after prolonged withdrawal from some forms of drug-conditioned training. Inhibition or removal of these receptors dampens drug-seeking behavior. However, it remains unclear which contextual variables associated with drug use govern CP-AMPAR expression and whether these receptors exhibit the long-term persistence expected of a reliable addiction-related biomarker. Here, we compared three fentanyl-conditioning settings in mice: conditioned place preference (CPP), open-field (OF), and home-cage (HC). Contrary to the HC condition, synaptic CP-AMPAR recruitment was observed in the NAc shell of both CPP and OF groups, which involved exposure to an unfamiliar context following drug administration. CPP and OF training produced comparable increases in CP-AMPAR expression, while CPP mice displayed heightened fentanyl-seeking behavior and OF mice showed locomotor sensitization. After more than 100 days of withdrawal, neither CP-AMPAR expression nor fentanyl-seeking behavior was detected following context re-exposure alone. However, fentanyl re-exposure reinstated drug-seeking behavior despite the absence of CP-AMPAR recruitment, which re-emerged only hours after drug administration. These findings indicate that contextual unfamiliarity drives CP-AMPAR recruitment after drug conditioning and suggest that these receptors transiently govern relapse in consecutive synaptic adaptations during withdrawal. The habituation or avoidance of contextual novelty during opioid treatments may therefore represent a previously underappreciated strategy to limit the recruitment of these maladaptive plasticity mechanisms and reduce relapse risk.

## INTRODUCTION

Recurrent episodes of drug relapse are a major challenge in treating substance use disorder (SUD) [1]. Drug relapse occurs even after prolonged abstinence and is triggered by contextual cues linked to prior drug experiences (conditioned stimuli), stress, or direct drug re-exposure (unconditioned stimuli) [2]. The persistent character of relapse vulnerability suggests that this propensity is driven by enduring, drug-induced adaptations within neural circuits governing motivated behavior. The brain region implicated in this process is the nucleus accumbens (NAc) [3,4], a central hub of reward processing and a critical gateway for the expression of addiction-related behaviors [5,6]. A pathological marker of plasticity evoked by repeated drug exposure in NAc medium spiny neurons (MSNs) is the persistent synaptic expression of Ca^2+^-permeable AMPA receptors (CP-AMPARs) following forced withdrawal [7,8]. These receptors feature distinct properties compared to other AMPAR subunit assemblies, such as the ability to conduct Ca^2+^, a second messenger, the exhibition of higher single-channel conductance, and inward rectification, characteristics that may amplify the responsiveness of MSNs to excitatory cortical and limbic inputs eliciting drug-seeking behaviors [9,10]. Importantly, pharmacological blockade or removal by optogenetic stimulation of CP-AMPARs in the NAc has been shown to significantly reduce drug-seeking responses in a rat self-administration procedure, indicating their functional role in relapse vulnerability [7,11]. Despite their ability to suppress CP-AMPAR activity, the effects of drug inhibitors are transient and do not permanently remove receptors, thereby hindering successful clinical translation of such approaches. Moreover, although these receptors have been regarded as a promising pathological hallmark of addiction-related plasticity, their long-term endurance after withdrawal has not been formally addressed. Understanding whether these receptors constitute a stable synaptic substrate and preventing their recruitment in the first place may prove valuable in mitigating relapse risk and reducing the likelihood of developing SUDs. We then sought to identify the specific conditions under which repeated drug exposure fails to induce CP-AMPAR expression and whether these receptors persist following long-term withdrawal.

It remains unresolved which contextual and procedural variables govern the incorporation of CP-AMPARs into the NAc following withdrawal. Initial studies reported that their expression emerged when drug exposure occurred through operant conditioning, where novel contextual cues were associated with the self-inflicted drug experience in a contingent administration procedure [7,11]. Subsequent work, however, independently reported CP-AMPAR upregulation following repeated experimenter-delivered drug injections [8,12], indicating that self-administration is not required. A key, but unaddressed variable in these experiments, is the context in which drug conditioning occurred, as the use of an open-field test and exposure to its arena may form a contingency between contextual cues and drug exposure. Yet across studies, contextual and drug-administration contingencies were not discriminated, and both were referred to simply as ‘contingent’ or ‘noncontingent’. When cocaine was administered in the animal’s home-cage with no exposure to a novel/unfamiliar environment (reported as noncontingent), CP-AMPARs did not accumulate after prolonged withdrawal [12]. Similarly, daily habituation to an open-field before cocaine injections (also reported as noncontingent) prevented CP-AMPAR expression [13]. In contrast, when mice were habituated to the test arena days before conditioning but not before each injection (also reported as noncontingent), CP-AMPARs were expressed [14]. The use of the terms ‘contingent’ and ‘noncontingent’ in these studies referred exclusively to the contingency of drug administration and undervalued the contingency of context. The necessary features of contextual contingencies for the induction of CP-AMPARs in the NAc thus remain insufficiently understood. CP-AMPARs have been linked to incubation of craving [7]. Since incubation increases during the first 2 months following withdrawal before diminishing after 6 months [15,16], it is assumed that CP-AMPAR expression prevails as well beyond the typically measured time points of 20–40 days. While these findings have suggested that CP-AMPAR expression accompanies relapse responses during early withdrawal, they are insufficient to determine whether receptor recruitment and its functional association with relapse risk persist across extended periods of abstinence. Consequently, it remains unclear whether CP-AMPAR expression reflects a long-term marker of relapse vulnerability or a more transient form of drug-induced plasticity.

Here, we investigated the role of contextual novelty/unfamiliarity in CP-AMPAR accumulation within the NAc shell following forced drug withdrawal, using three distinct experimental settings. To circumvent the methodological conflicts presented by habituation-dependent noncontingent context paradigms, we employed home-cage injections as our noncontingent setting condition, ensuring drug exposure occurred without pairing to an unfamiliar context. This approach, while precluding the assessment of locomotor sensitization, allowed for a rigorous isolation of contextual novelty effects. The other two procedures, the open-field and the conditioned place preference, provided contexts where drug administration was accompanied by exposure to unfamiliar cues. We selected fentanyl as the test drug, given the potential implications that contextual unfamiliarity may have for relapse vulnerability in clinical treatments with opioids.

We then carried out long-term conditioned place preference tests to assess whether CP-AMPAR expression persists across longer withdrawal intervals and remains associated with relapse-related behaviors. Following extended abstinence, mice underwent either contextual re-exposure or fentanyl re-exposure before electrophysiological assessment of relative CP-AMPAR levels. This approach allowed us to determine whether CP-AMPARs represent a lasting synaptic adaptation linked to relapse vulnerability or a transiently regulated mechanism of addiction-related plasticity.

## MATERIALS AND METHODS

### Mice

Male and female C57BL/6J background mice (Charles River), aged 4-5 weeks at the start of the experiments, were housed in groups of 4 (males) or 5 (females) under a 12-hour light/dark cycle. All experiments were conducted during the light phase. Procedures were approved by all respective government authorities (IACUC, University of Pittsburgh).

### Conditioned Place Preference (CPP)

Mice were habituated to the testing room and the experimenter for 5 days, during which they received two intraperitoneal (i.p.) sham injections. Conditioned place preference (CPP) was assessed using an apparatus consisting of two conditioning chambers (15 x 15 cm each) separated by a neutral compartment with guillotine doors. The chambers were differentiated by distinct sensory cues: olfactory (a piece of tissue with a drop of fresh linen or cinnamon essential oil, sealed inside a closed Eppendorf tube), tactile (circular or striped linear grooves on the floor), and visual (one chamber with thick vertical black-and-white stripes, the other with small black dots on a white background). On the day prior to conditioning (D0), mice were placed in the neutral zone and allowed free exploration of both chambers for 20 minutes to establish a baseline (BL) preference. Time spent in each chamber was recorded, and baseline preference for the striped chamber was calculated as: Stripe Time / (Stripe Time + Dot Time) x 100. Animals displaying strong initial bias (>70% preference for either chamber) were excluded from further testing. During the 10-day conditioning phase, each mouse was assigned a fentanyl-paired chamber (CS^+^). Mice received alternating i.p. injections of fentanyl (0.2 mg/kg) or saline, and were immediately placed in the corresponding chamber for 20 minutes. Following conditioning, mice underwent a 20-day forced withdrawal period. Conditioned preference was evaluated twice, with respect to the CS^+^ chamber for the fentanyl group and the stripe chamber for the saline group: once on the first day of withdrawal (WD1, Post-test I) and again on the final day (WD20, Post-test II). During both post-tests, mice were given free access to both chambers without drug or saline administration. After Post-test II, animals were sacrificed within 10 minutes for electrophysiological analysis.

### Open-field (OF)

For the open-field (OF) paradigm, mice were habituated to the experimental room and the experimenter for five consecutive days, following the same general schedule described in the CPP methods subsection. The open-field apparatus consisted of a square arena (31 x 20 x 31 cm) with white walls and floor and an open top. The conditioning schedule lasted five days. Each day, mice received a single intraperitoneal (i.p.) injection of fentanyl (0.2 mg/kg) or saline, according to group assignment, and were immediately placed in the center of the arena. Each session lasted 20 minutes, during which the animals were video recorded for behavioral analysis. Locomotor activity was quantified using the first 10 min of each recording. Total distance traveled and average speed were estimated with the AnimalTracker plugin in Fiji [38,39] and used to assess locomotor sensitization to fentanyl exposure across training sessions. Following completion of the training period, mice underwent a 20-day forced withdrawal phase. On the final day (WD20), animals were euthanized for electrophysiological recordings.

### Home-cage (HC)

For the home-cage (HC) setting, mice were habituated to experimenter handling for five consecutive days, during which they received two i.p. saline injections. During conditioning, mice received 0.2 mg/kg fentanyl daily for five days and were immediately returned to their standard housing cages (7 x 12 x 27 cm) after each administration. Following completion of the conditioning schedule, animals underwent a 20-day forced withdrawal period. On WD20, mice were sacrificed for electrophysiology.

### Long-term withdrawal tests

To assess the long-term persistence of CP-AMPARs following extended withdrawal, mice underwent the same conditioned place preference (CPP) training procedure described above. The only modification to the original schedule was that the second preference test (Post-test II) was conducted after more than 100 days of withdrawal (WD103-WD114) rather than on WD20. Five experimental groups were included, consisting of four fentanyl-conditioned groups and one saline-conditioned group. The Re-exposure group underwent Post-test II on WD103-WD114 and was sacrificed for electrophysiological assessment immediately thereafter. The Fentanyl group received an intraperitoneal fentanyl injection (0.2 mg/kg) immediately before Post-test II and was sacrificed immediately following behavioral testing. The Fentanyl +6h group received the same fentanyl injection and underwent Post-test II, but electrophysiological recordings were performed 6 hours later. An electrophysiological Control group did not undergo CPP re-exposure or fentanyl re-exposure; mice were sacrificed directly for electrophysiological assessment on their corresponding withdrawal day. Saline-conditioned animals underwent the same procedure as the Re-exposure group and served as behavioral and electrophysiological controls.

### Nucleus accumbens coronal slice preparation

Mice were deeply anesthetized with isoflurane and subsequently decapitated. Coronal brain slices (300 µm) were prepared using a vibratome in an ice-cold NMDG-based cutting solution (135 mM N-methyl-D-glucamine, 10 mM D-glucose, 1 mM KCl, 1.5 mM MgCl_2_, 0.5 mM CaCl_2_, 1.2 mM KH_2_PO_4_, 20 mM choline bicarbonate). Slices were then transferred to oxygenated aCSF solution (119 mM NaCl, 26 mM NaHCO_3_, 20 mM D-glucose, 2.5 mM KCl, 1 mM NaH_2_PO_4_, 1.3 mM MgSO_4_ • 7H_2_O, 2.5 mM CaCl_2_) and incubated at 34°C for 20 minutes. After incubation, slices were maintained at room temperature in continuously oxygenated aCSF for at least 1 hour before electrophysiological recordings.

### Electrophysiology

Standard whole-cell recordings in voltage-clamp mode were carried out at 30 ± 2°C in a recording chamber continuously perfused with ACSF (2 mL/min). Medium spiny neurons of the NAc shell were visually identified with infrared-differential interference contrast microscopy. Borosilicate glass pipettes (2-5 MΩ) were filled with Cs-based internal solution (120 mM Cs-gluconate, 20 mM HEPES, 0.4 mM EGTA, 2.8 mM NaCl, 5 mM TEA-Cl, 4 mM Mg-ATP, 0.3 mM Na-GTP, 100 µM spermine, pH 7.2). Electrical stimulation was provided through tungsten metal electrodes. Recordings were acquired using custom algorithms in Igor Pro (Wavemetrics) with an ELC-03XS amplifier (NPI) and digitized at 10 kHz with an ITC-18 interface (HEKA). Data were filtered at 3 kHz, and series and input resistances were monitored throughout recording sessions. GABA-A receptor-mediated transmission was pharmacologically inhibited by applying 100 µM picrotoxin to the perfusate across all recordings. To isolate AMPAR responses, NMDAR currents were blocked by application of 50 µM D-APV to the aCSF. EPSCs were recorded at holding potentials of -60 mV and +40 mV at 0.1 Hz. The CP-AMPAR rectification index (RI) was calculated as: RI = mean EPSC amplitude at -60 mV/mean EPSC amplitude at +40 mV. Each averaged EPSC trace reflects ∼30 stimulation sweeps at each holding potential.

### Data analysis

Results are reported as mean ± standard deviation (SD). Behavioral CPP data were analyzed using repeated-measures one-way ANOVA. Open-field locomotor activity data (total distance and average speed) and two-group electrophysiological comparisons were analyzed using unpaired two-tailed Student’s t-tests. Open-field sensitization data were analyzed using a mixed-effects model (REML) with Tukey’s multiple comparisons test. Electrophysiological data involving more than two groups were analyzed using ordinary one-way ANOVA. Statistical significance is indicated as *p<0.05, **p<0.01, ***p<0.001, ****p<0.0001, and ns>0.05. Electrophysiological analyses were performed using animal-based statistics: data from multiple cells recorded in the same animal were averaged to yield a single value per animal, thereby avoiding pseudoreplication. Open-field behavioral analyses were likewise performed using animal-based statistics, with measurements from multiple recording sessions averaged to obtain a single value per animal before statistical testing. Sample sizes are reported as *n/m* for electrophysiological experiments, where *n* denotes the number of animals and *m* the number of recorded cells.

## RESULTS

### CPP and Open-field training similarly elevated synaptic CP-AMPARs in the NAc shell after fentanyl conditioning

To test the impact of contextual novelty/unfamiliarity during drug conditioning on CP-AMPAR acquisition, we first trained WT mice in a conditioned place preference (CPP) apparatus to associate one of the cued chambers with fentanyl exposure (CS^+^). Both cocaine and fentanyl have been reported to induce CP-AMPARs in NAc MSNs after withdrawal following CPP conditioning [10,12]. We placed mice in the CS^+^ chamber immediately after receiving a 0.2 mg/kg injection of fentanyl once every other day, while the alternate chamber was paired with saline, the conditioned control stimulus (CS^-^). To evaluate whether a preference for the CS^+^ chamber was established, and whether this cue-drug memory was retained after withdrawal, mice were first allowed to freely explore the open apparatus on day 0 of training (D0) to set a baseline value (BL), then on withdrawal day 1 (D11/WD1; one day after training completion) and withdrawal day 20 (WD20; the day of electrophysiological analysis) (Fig. 1a). Since mice were exposed to the apparatus on D0, the CPP context at the start of the training schedule was no longer novel but rather unfamiliar, as mice were also not habituated to the apparatus on the day of training.

**Figure 1.**
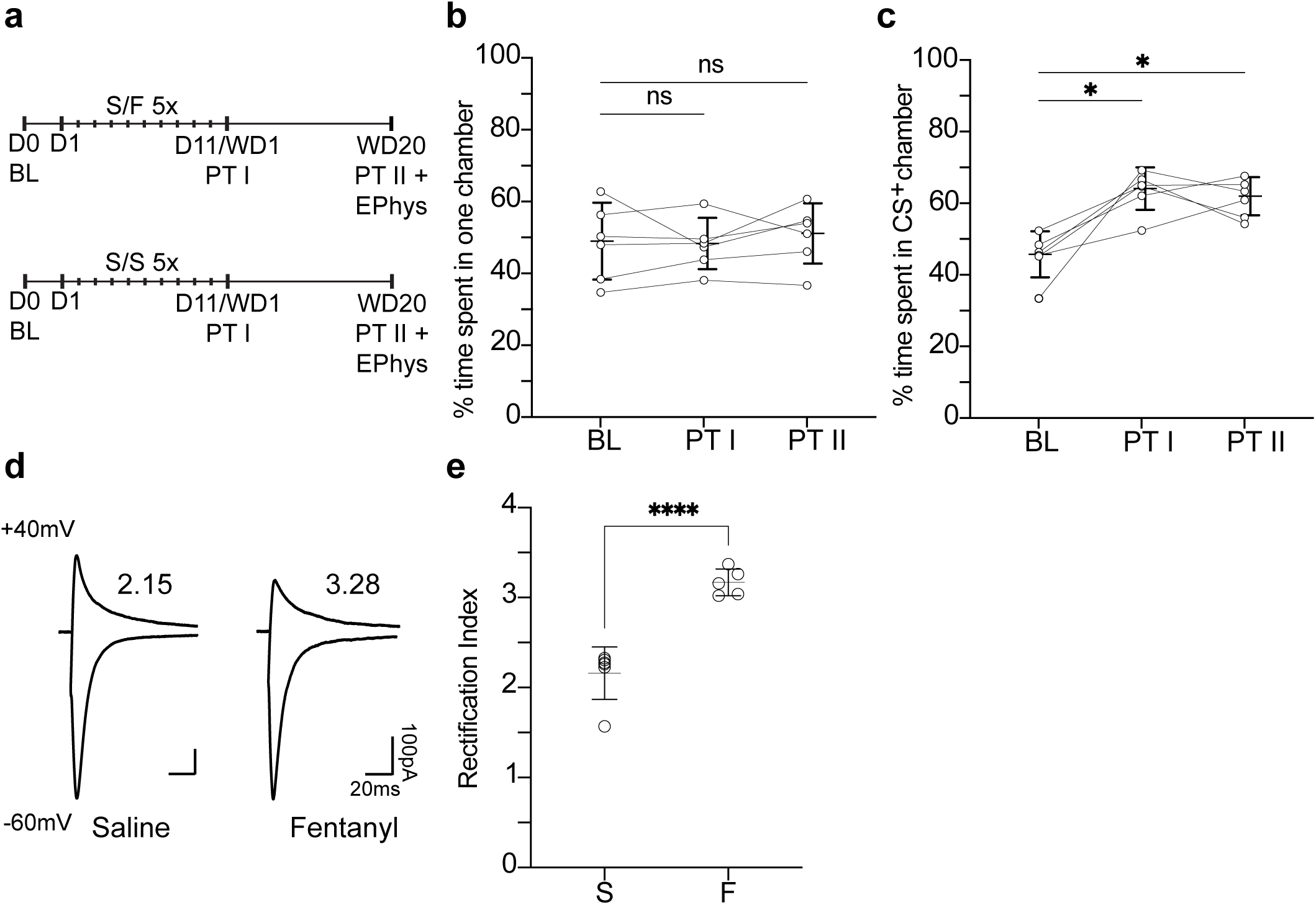
CPP training elevated synaptic CP-AMPARs in the NAc shell after fentanyl conditioning. **a** Diagram illustrating behavioral training and electrophysiology schedule for **CPP** groups (D#–day #, BL–baseline, PT #–Post-Test #, WD#–withdrawal day #). **b, c** CPP was measured at three timepoints, BL (D0), PT I (WD1), PT II (WD20), and calculated based on time spent on CS^+^ (fentanyl group) or striped (control group) chamber. Statistical significance is indicated as *p<0.05, ****p<0.0001, and ns>0.05. **d** Sample average AMPA EPSCs evoked at +40 and -60 mV in NAc shell MSNs from Saline and Fentanyl groups, with the corresponding RI value for each trace. **e** Rectification index of AMPAR EPSCs (saline WD20, n/m = 6/20; fentanyl WD20, n/m = 5/16) evoked at +40 and -60 mV in NAc shell MSNs of WT mice. Smaller dots in **e** represent individual mice. n/m = number of animals/number of cells recorded.

Mice paired with the CS^+^ chamber developed a fentanyl-CPP preference by WD1 compared to the D0 baseline assessment, indicating the formation of an associative link between chamber cues and fentanyl exposure (Fig. 1c; for Pre-test: 45.18 ± 6.34; for Post-test I: 63.33 ± 5.84; n = 6; Pre-test vs Post-test I, *p<0.05*). On WD20, the preference for the CS^+^ chamber remained, reflected by persistent fentanyl-seeking behavior comparable to that observed on WD1 (Fig. 1c; for Post-test II: 61.20 ± 5.24, n = 6; Pre-test vs Post-test II, *p<0.05*). Control mice had no chamber preference on either WD1 or WD20 (Fig. 1b; for Pre-test: 48.42 ± 10.59; for Post-test I: 47.77 ± 7.05; for Post-test II: 50.55 ± 8.30; n = 6; Pre-test vs Post-test I, *p=0.9686*; Pre-test vs Post-test II, *p=0.7411*), demonstrating that the arrangement of the CPP apparatus alone was insufficient to generate a side preference.

On WD20, the rectification index of AMPAR EPSCs was measured to assess the presence of CP-AMPARs in both groups through whole-cell patch clamp recordings of NAc shell MSNs and calculated as the ratio of -60 mV EPSC amplitude to the +40 mV EPSC amplitude (Fig. 1a). The rectification index was elevated for the CS^+^ mice compared to that of CS^-^ mice, indicating the expression of CP-AMPARs (Fig. 1e; for saline: 2.16 ± 0.29, n/m = 6/20; for fentanyl: 3.15 ± 0.15, n/m = 5/16; Saline vs Fentanyl, *p<0.0001*). These findings affirm previous reports and set the positive control for our study [10].

In a second experimental group, we examined whether CP-AMPARs are expressed in MSNs of the NAc shell following withdrawal from a fentanyl-open-field (OF) procedure. For 5 consecutive days, mice received 0.2 mg/kg fentanyl through i.p. injections shortly before being placed in the OF arena for 20 minutes. A control group received saline injections instead. Each session was video recorded to assess locomotor responses throughout the conditioning period. Fentanyl-treated mice exhibited greater locomotor activity compared to saline controls, reflected by increased total distance traveled and higher average speed during OF sessions (Fig. 2b-c; for total distance in centimeters, saline: 1236.61 ± 328.9, n = 5; fentanyl: 7646.13 ± 2605; n = 5; Saline vs Fentanyl, *p<0.001*. For average speed in meters per second, saline: 2.16 ± 0.52, n = 5; fentanyl: 12.74 ± 4.34, n = 5; Saline vs Fentanyl, *p<0.001*), and exhibited locomotor sensitization across repeated OF sessions, with average speed significantly increased on days 2-5 compared with day 1 (Fig. 2d; for average speed in meters per second, fentanyl D1: 12.311 ± 3.328, n = 5; D2: 15.060 ± 2.171, n = 4; D3: 14.100 ± 4.001, n = 4; D4: 15.302 ± 2.906, n = 4; D5: 15.324 ± 4.780, n = 4; D2 vs D1, *p<0.05*; D3 vs D1, *p<0.05*; D4 vs D1, *p<0.05*; D5 vs D1, *p<0.01*; no significant differences were observed across days in saline-treated mice). Following the completion of the training schedule, mice were subjected to forced withdrawal for 20 days. On WD20, the AMPAR rectification indices were measured (Fig. 2a). AMPAR EPSCs of mice that received fentanyl injections during OF training exhibited higher inward rectification than saline-injected mice (Fig. 2f; for saline: 2.38 ± 0.20, n/m = 4/13; for fentanyl: 3.20 ± 0.11, n/m = 4/16; Saline vs Fentanyl, *p<0.001*).

**Figure 2.**
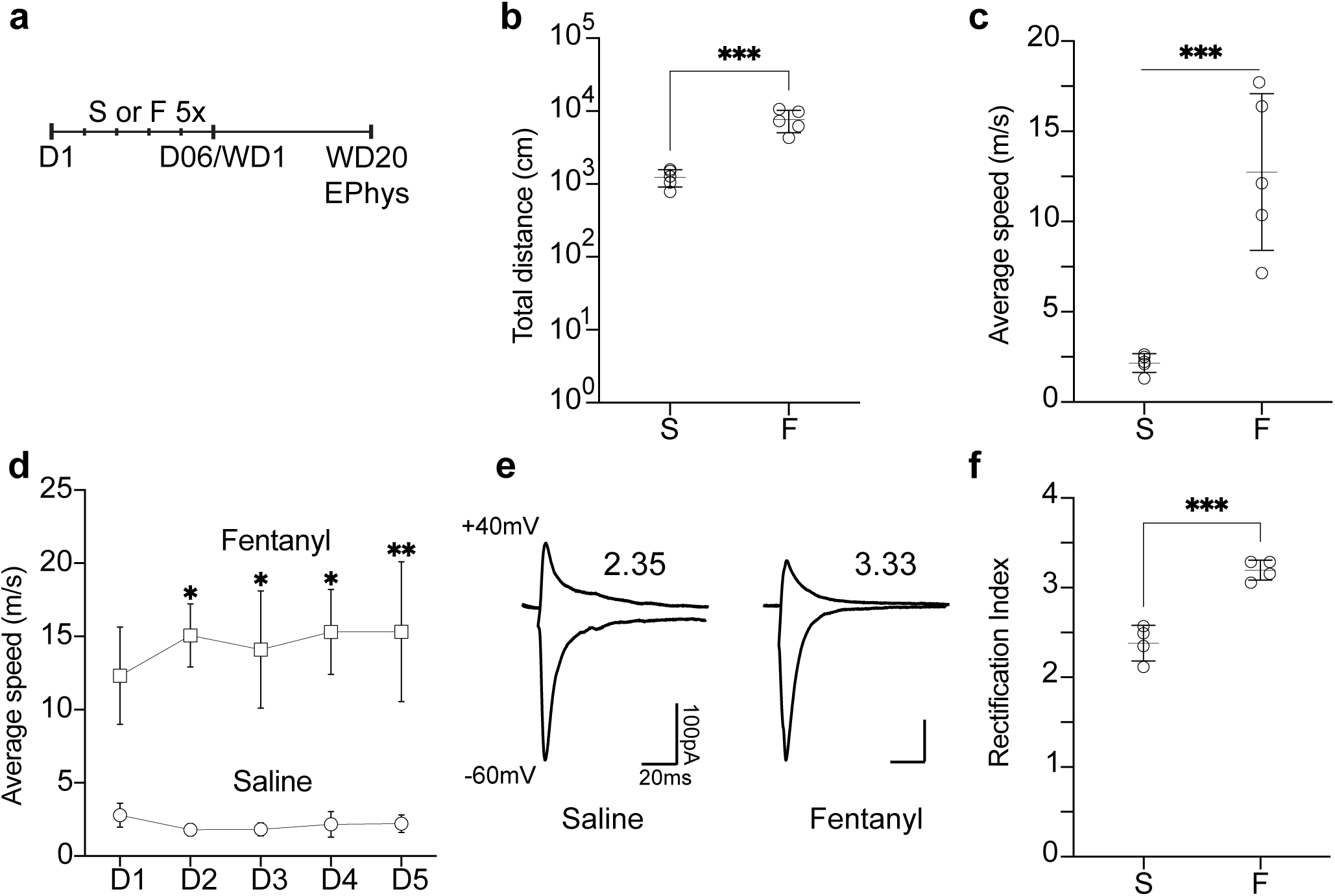
Open-field training elevated synaptic CP-AMPARs in the NAc shell after fentanyl conditioning. **a** Diagram illustrating behavioral training and electrophysiology schedule for **OF** groups (D#–day #, WD#–withdrawal day #). **b, c, d** Locomotor activity and sensitization during OF training, assessed from video recordings by total distance traveled (**b**), average speed (**c**), and average speed across training days (**d**). Data are reported as animal averages (n = 5 per group for total distance and average speed; n varied across days for average speed). Statistical significance is indicated as *p<0.05, **p<0.01, ***p<0.001. **e** Sample average AMPA EPSCs evoked at +40 and -60 mV in NAc shell MSNs from Saline and Fentanyl groups, with the corresponding RI value for each trace. **f** Rectification index of AMPAR EPSCs (saline WD20, n/m = 4/13; fentanyl WD20, n/m = 4/16) evoked at +40 and -60 mV in NAc shell MSNs of WT mice. Smaller dots in **f** represent individual mice. n/m = number of animals/number of cells recorded. Statistical significance is indicated as ***p<0.001.

### Fentanyl training in the home-cage failed to induce CP-AMPAR expression in the NAc shell

In a final setting evaluating contextual variables, we examined whether fentanyl administration generally induces CP-AMPAR induction after withdrawal independent of contextual contingencies. Previous studies with cocaine have reported conflicting results under different noncontingent administration conditions [8,12–14]. As for cocaine, we used the home-cage setting for the fentanyl administration under both noncontingent administration and context conditions [12]. In this setting, mice received 0.2 mg/kg fentanyl injections in a contextually familiar setting as we returned them directly to their home-cages (HC) after the drug administration. Conditioning in the HC paradigm followed the same schedule as that in the OF groups. At WD20, the AMPAR rectification index did not differ between the fentanyl- and saline-treated HC mice (Fig. 3c; for saline: 2.14 ± 0.23, n/m = 4/14; for fentanyl: 2.39 ± 0.21, n/m = 5/16; Saline vs Fentanyl, *p=0.1842*), and was similar to the control groups from the OF and CPP settings (OF Saline vs HC Fentanyl, *p=0.9709*; CPP Saline vs HC Fentanyl, *p=0.1687*), indicating that repeated fentanyl injections alone are insufficient to drive CP-AMPAR upregulation. Therefore, repeated drug administration required a concomitant exposure to contextual novelty/unfamiliarity during drug conditioning (i.e., a contextual contingency) to induce the expression of CP-AMPARs in the NAc shell MSNs.

**Figure 3.**
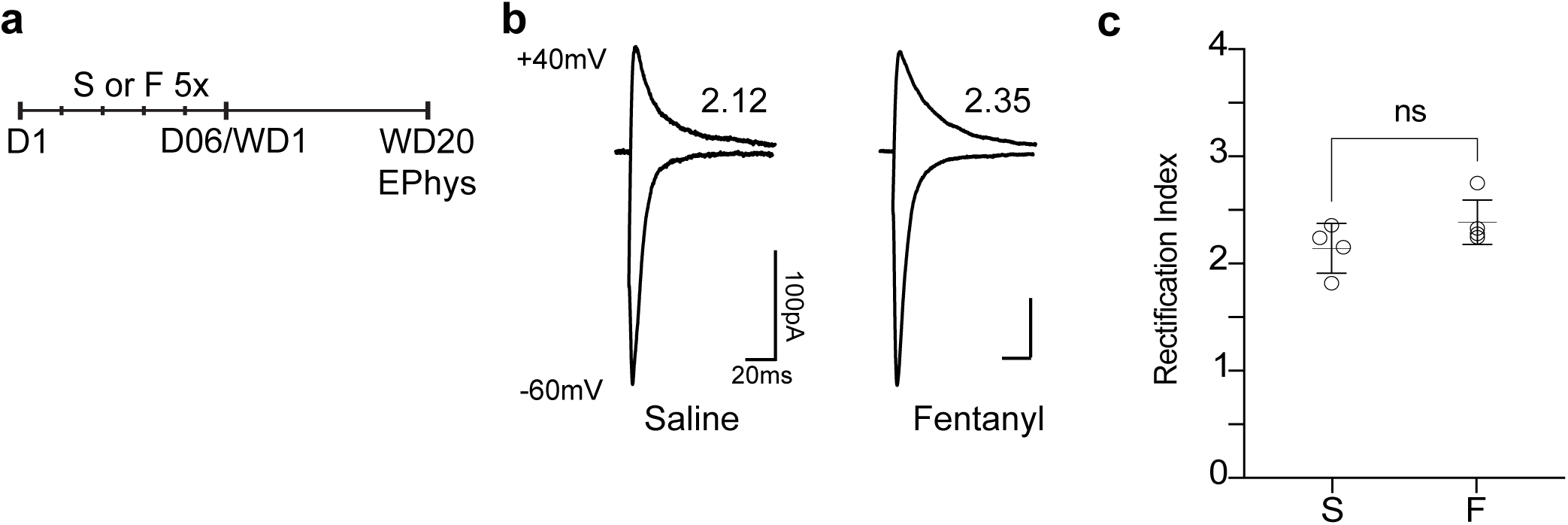
Fentanyl training in the home-cage failed to induce CP-AMPAR expression in the NAc shell. **a** Diagram illustrating behavioral training and electrophysiology schedule for HC groups (D#–day #, WD#–withdrawal day #). **b** Sample average AMPA EPSCs evoked at +40 and -60 mV in NAc shell MSNs from Saline and Fentanyl groups, with the corresponding RI value for each trace. **c** Rectification index of AMPAR EPSCs (saline WD20, n/m = 4/14; fentanyl WD20, n/m = 5/16) evoked at +40 and -60 mV in NAc shell MSNs of WT mice. Smaller dots in **c** represent individual mice. n/m = number of animals/number of cells recorded. Statistical significance is indicated as ns>0.05.

### CP-AMPAR expression was absent following long-term withdrawal and re-emerged only after fentanyl re-exposure

To address the limitations of relatively short withdrawal periods found in previous studies examining CP-AMPAR expression, we next asked whether these receptors remain incorporated after more than 100 days of abstinence and continue to accompany relapse-related responses. Following long-term withdrawal (WD>100), CPP re-exposure alone failed to elicit a preference for the CS^+^ chamber, indicating that the conditioned drug memory was no longer sufficient to drive fentanyl-seeking behavior (Fig. 4b; for Pre-test: 46.05 ± 3.00; for Post-test I: 65.10 ± 5.25; for Post-test II: 49.07 ± 6.95; n = 5; Pre-test vs Post-test II, *p=0.7553*; Post-test I vs Post-test II, *p=0.0631*). In contrast, fentanyl re-exposure before testing, reinstated CPP (Fig. 4c; for Pre-test: 46.36 ± 3.22; for Post-test I: 61.87 ± 4.53; for Post-test II: 67.26 ± 17.50; n = 8; Pre-test vs Post-test II, *p<0.05*; Post-test I vs Post-test II, *p=0.6340*), indicating that the memory trace was indeed present and remained susceptible to reactivation by drug exposure.

**Figure 4.**
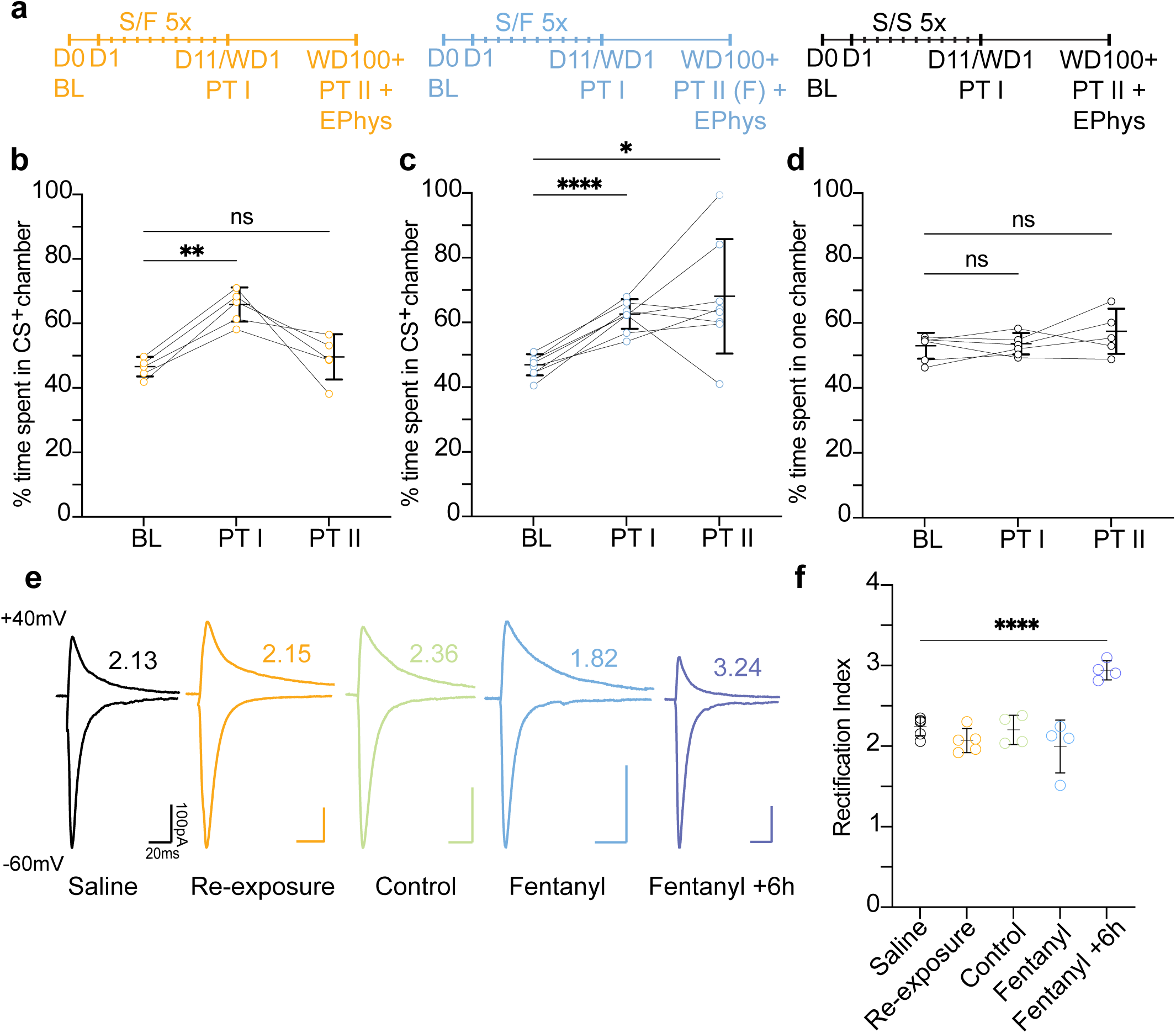
CP-AMPAR expression was absent following long-term withdrawal and re-emerged only after fentanyl re-exposure. **a** Diagram illustrating behavioral training, long-term withdrawal, CPP re-exposure, and electrophysiology schedules for **Re-exposure**, **Fentanyl**, and **Saline** groups (D#–day #, BL–baseline, PT #–Post-Test #, WD#–withdrawal day #). **b-d** CPP was measured at three timepoints, BL (D0), PT I (WD1), and PT II (>WD100), and calculated based on time spent on the CS+ chamber for **Re-exposure** (**b**) and **Fentanyl** (**c**) groups, or the striped chamber for the **Saline** group (**d**). **e** Sample average AMPA EPSCs evoked at +40 and -60 mV in NAc shell MSNs from Saline, Re-exposure, Control, Fentanyl, and Fentanyl +6h groups, with the corresponding RI value for each trace. **f** Rectification index of AMPAR EPSCs (saline, n/m = 6/15; re-exposure, n/m = 5/17; control, n/m = 4/11; fentanyl, n/m = 4/12; fentanyl +6h, n/m = 4/14) evoked at +40 and -60 mV in NAc shell MSNs of WT mice. Smaller dots in **f** represent individual mice. n/m = number of animals/number of cells recorded. Statistical significance is indicated as *p<0.05, **p<0.01, ****p<0.0001, and ns>0.05.

We next assessed CP-AMPAR expression in NAc shell MSNs under each condition. Neither the electrophysiological control group nor mice exposed only to the CPP apparatus exhibited elevated rectification indices relative to saline controls (Fig. 4f; for saline: 2.25 ± 0.11, n/m = 6/15; for control: 2.20 ± 0.15, n/m = 4/11; for re-exposure, 2.07 ± 0.15, n/m = 5/17; Saline vs Control, *p=0.9951*; Saline vs Re-exposure, *p=0.5310*), indicating that CP-AMPAR expression was no longer detectable after long-term withdrawal. Notably, CP-AMPARs also remained absent immediately following fentanyl-induced reinstatement despite the recovery of drug-seeking behavior (Fig. 4f; for fentanyl, 2.00 ± 0.33, n/m = 4/12; Saline vs Fentanyl, *p=0.2613*). However, when electrophysiological recordings were performed 6 hours after fentanyl re-exposure, rectification indices were significantly increased (Fig. 4f; for fentanyl +6h, 2.94 ± 0.12, n/m = 4/14; Saline vs Fentanyl +6h, *p<0.0001*), indicating renewed CP-AMPAR recruitment. These findings demonstrate that CP-AMPAR expression is not maintained throughout extended withdrawal and is absent at the time of memory reinstatement, but can be rapidly re-engaged following re-exposure to the unconditioned stimuli elicited by drug use.

## DISCUSSION

We found that noncontingent drug administration was sufficient to induce CP-AMPAR expression in NAc shell MSNs following drug withdrawal as long as contextual contingencies were present. In other words, this form of maladaptive synaptic plasticity was prevented when drug exposure occurred in the absence of novel/unfamiliar contextual cues. These findings may have translational implications for the contextual settings in which opioid therapies occur to reduce relapse risk and drug craving, which have been implicated by NAc CP-AMPAR presence. The NAc shell CP-AMPARs were also expressed for several weeks after fentanyl withdrawal but eventually were removed to only reappear with fentanyl re-exposure after long-term withdrawal, suggesting specific consecutive synaptic adaptations during withdrawal.

Experimenter-delivered fentanyl during both CPP and open-field conditioning induced CP-AMPAR expression in NAc shell MSNs after 20 days of withdrawal. In contrast, fentanyl administered in the home-cage did not induce synaptic CP-AMPAR expression. These results extend our previous findings for CPP and home-cage cocaine administration to an opioid model [12]. They are also consistent with evidence that cocaine exposure in an open-field test induced CP-AMPARs [14], yet appear at odds with another study reporting no CP-AMPAR recruitment under a seemingly comparable open-field procedure [13]. This discrepancy cannot be explained by differences in drug identity or NAc subregion, as the absence of CP-AMPARs in the cocaine open-field procedure was observed in both NAc shell and core MSNs [13], whereas the presence of CP-AMPARs after cocaine and fentanyl was specifically investigated in the NAc shell [14] (Figs. 1e, 2f, 3c, 4f). Methodological differences are then likely responsible for the divergent outcomes. We and Terrier et al. (2016) administered the drug without daily habituation to the open-field arena and observed the induction of synaptic CP-AMPARs. In contrast, McCutcheon et al. (2011) habituated the animals to the arena each day before i.p. cocaine injections. Thus, with a habituation schedule as that applied by McCutcheon et al. (2011), rodents might acquire a state of context familiarity, functionally replicating a home-cage setting condition. This implies that CP-AMPAR expression induced by the contingency between contextual cues and drug experience is sensitive to the length of prior exposure to that context (i.e., familiarity with the context). Only unfamiliar settings, such as the CPP apparatus, the open-field test arena without prior habituation, or the operant conditioning chamber, appear to induce CP-AMPAR expression in NAc shell MSNs. Notably, habituation carried out over a few days before the onset of cocaine conditioning, but not on the days of drug administration themselves, does not sufficiently reduce the unfamiliarity effect of the context to prevent CP-AMPAR expression [14].

Induction of CP-AMPAR expression was also reported in another experimenter-administered cocaine study [8]. However, the provided procedural details were insufficient to determine the precise contextual contingencies under which drug exposure occurred. Notably, that cohort included younger mice than the other studies, raising the possibility that their young age made them more susceptible to CP-AMPAR expression.

The NAc core and shell respond differently to the extent of operant conditioning in terms of inducing synaptic CP-AMPARs and dendritic spine density [9,17]. Long-access cocaine training sessions induce the expression of CP-AMPARs in both the NAc core and shell after withdrawal [7,13], whereas short-access cocaine training sessions do not induce CP-AMPARs in the NAc core [9]. However, during withdrawal, short-access cocaine induces CP-AMPARs in the NAc shell, which govern the incubation of craving [11]. Either short- or long-access to cocaine causes an increase in spine density, an effect that persists for over a month [17]. However, spine increase is more pronounced in the NAc core selectively after long-access to cocaine. Thus, the NAc core may express additional maladaptive plasticity mechanisms with extended drug access.

For CPP, it is not yet known whether CP-AMPAR induction is restricted to the NAc shell or also occurs in core MSNs [10,12,18] (Figs. 1e, 4f). Notably, incubation of craving has also been reported for morphine-CPP, hence with passive drug administration, when low training doses were used, indicated by a higher preference score at WD14 relative to WD1 [19], while higher doses yield a similar preference score at both withdrawal time points. However, it was not tested whether CP-AMPAR expression underlies the higher CS^+^ chamber preference.

In contrast, this was shown for the reinstatement of drug seeking [10]. A low dose of cocaine or fentanyl can reinstate drug seeking only when CP-AMPARs are present, whereas in their absence, a higher dose is required. Together, these results support a role for CP-AMPARs in both the NAc core and shell subregions in regulating the intensity of drug-seeking behavior and the threshold at which contextual cues can trigger cue-drug memory retrieval and reinstatement of seeking. These two regulatory effects correspond to craving and relapse vulnerability, respectively, and are central features of SUD symptomatology. Collectively, these studies provide evidence for a model in which contextual contingencies with drugs of abuse induce CP-AMPARs in the NAc shell. Conversely, the NAc core expresses additional plastic changes with long-term access to drug intake, likely facilitating the escalation toward compulsive drug use [20].

Our findings also provide new and key insight into the long-term behavior of CP-AMPARs following extended periods of withdrawal and complement and extend, rather than contradict, previous studies on their role in relapse vulnerability and drug-generated memories. Re-exposure to conditioned stimuli alone was insufficient to elicit drug-seeking behavior after more than 100 days of withdrawal, indicating that the previously established cue-drug association was no longer readily retrievable under these conditions, but rather required a stronger trigger to become reactivated. Fentanyl re-exposure, one such trigger, reinstated drug-seeking behavior. This occurred without the immediate presence of CP-AMPARs, which resurfaced only several hours after the fentanyl injection. These observations suggest that CP-AMPAR expression is not constitutively maintained throughout long-term abstinence and that this associative link with relapse behavior may be temporally restricted. While previous research demonstrated a clear relationship between CP-AMPARs and relapse risk during earlier time periods of withdrawal, our results heavily constrain and show that this relationship becomes less direct over longer timescales.

The absence of both CPP expression and CP-AMPAR recruitment following context re-exposure alone is further consistent with the view that CP-AMPARs regulate the retrieval of drug-associated memories rather than their long-term storage. At the same time, the rapid re-emergence of these receptors following fentanyl re-exposure suggests that the cellular processes responsible for CP-AMPAR recruitment remain preserved long after withdrawal and are liable to be re-engaged. One candidate interpretation is that CP-AMPARs function as dynamically regulated facilitators of drug-memory expression, becoming involved when drug-induced memory traces have recently been activated or are particularly susceptible to strengthening or modification, rather than constituting the trace itself. Whether CP-AMPAR recruitment during early withdrawal is required for the consolidation and maturation process of drug-associated memories remains unknown. Future studies combining prolonged withdrawal paradigms like ours with pharmacological blockade of CP-AMPARs may clarify whether these receptors may constitute a critical gateway in shaping the enduring properties of these memories.

A period of 2-3 months after drug withdrawal also appears to more generally alter drug-associated behaviors. While increased cue induced drug seeking after drug-self-administration is observed 1 month after withdrawal [7,11,16], it is reduced to WD1 values after 2 or 3 months of withdrawal for heroin or cocaine self-administration [21,22]. It is unknown whether this decline also parallels the removal of CP-AMPARs in NAc MSNs.

Our results carry important implications for the prevention of opioid use disorder (OUD), and, more broadly, substance use disorders (SUDs) arising from medically prescribed drugs. Although treatment approaches for OUD remain a major focus in addressing the urgency of the opioid crisis, preventive strategies have yet to advance considerably beyond the regulation of prescription practices [23]. Our results indicate that the degree of habituation to the environment in which opioids are administered is a potential risk factor for relapse development. Contextual unfamiliarity during prescribed opioid use thus warrants attention in the design and implementation of clinical interventions, and a period of contextual habituation before drug administration appears adequate to attenuate this unfamiliarity effect, based on our results and the re-evaluation and comparison of previous works [13,14]. While such an approach may only be applicable to cases of accidental, iatrogenic OUD rather than deliberate misuse, such as with surgical patients [24], its relevance remains pronounced given the high abuse potential of opioids like fentanyl and the fact that addiction vulnerability cannot be fully prevented by adherence to medical guidelines alone [25,26]. The lack of CP-AMPARs post-withdrawal in a noncontingent setting condition can also partly explain why hospitalized patients receiving potent opioids for postoperative or palliative care may not develop addiction-like symptoms, as these medical contexts are generally familiar or can be readily habituated to.

Another relevant comparison emerging from our results is that between the fentanyl-treated OF and CPP groups. The increase in CP-AMPAR expression observed in the CPP setting does not seem to result from re-exposure to the conditioning context during Post-test II, since the OF group displayed similarly elevated rectification indices despite not undergoing any re-exposure procedure at WD20. The absence of a difference in the RI values between the two procedures suggests that the presence of contextual novelty/unfamiliarity, irrespective of its relative abundance or complexity, is sufficient to drive comparable levels of CP-AMPAR recruitment. One could have hypothesized that the CPP apparatus, being more densely populated with novel sensory cues than the OF arena, might yield a stronger rectification effect through a higher number of potential drug-cue associations. However, while a greater abundance of associations could conceivably strengthen the cue-drug memory resulting from conditioning, the post-withdrawal incorporation of CP-AMPARs may instead represent an all-or-none process, largely independent of further contextual variations beyond the simple, baseline unfamiliarity level.

The mechanistic details of the unfamiliarity effect in inducing CP-AMPAR expression under contextual contingencies need further exploration. It has been proposed that novelty perception is heterogeneous and can be categorized into two functionally distinct types, contextual (common or distinct) [27] and absolute novelty, both engaging the anterior hippocampus but differing in their neuromodulatory signatures [28]. While contextual novelty likely represents a form of novelty based on deviations from established memory schemas, recruiting the hippocampal-midbrain loop and eliciting dopaminergic input from the ventral tegmental area and noradrenergic input from the locus coeruleus [29–32], absolute novelty corresponds to the exposure to an entirely new context and it appears to involve a cholinergic-hippocampal pathway, which in turn potentially modulates hippocampal encoding through a reduced interference from prior representations [33,34]. Although the relationship between these parallel systems may not be strictly dichotomous and might present some overlap, their relative contributions may shift dynamically with experience. It is conceivable that the novelty of the OF and the novelty of the CPP environments initially engage an absolute novelty response (cholinergic), and, upon the CPP Pre-test and the CPP and OF repeated drug training exposure, transition toward dopaminergic and noradrenergic signaling. Such signals may underlie a neuromodulatory gating process in which novelty-driven arousal facilitates or enables subsequent CP-AMPAR acquisition following drug conditioning. While this needs further experimentation, it provides a plausible mechanistic substrate for the unfamiliarity effect associated with contextually contingent drug exposure and relapse vulnerability.

The insertion of CP-AMPARs into principal cells of the NAc after drug abstinence represents a promising form of synaptic plasticity contributing to the persistence of relapse episodes even long after the disappearance of withdrawal symptoms [7,10]. In the present study, we demonstrated that the acquisition of these receptors depends on repeated drug exposure within an unfamiliar context, and reinforced the idea that this effect is not dependent on a contingent administration procedure. The degree of acquaintance with the environment where drug intake occurs thus emerges as an important variable influencing the development of relapse vulnerability. We additionally showed that CP-AMPAR recruitment is not maintained after long-term withdrawal, but instead is dynamically re-engaged following renewed drug exposure, thereby suggesting that the temporal window over which this plasticity effect is expressed and associated with drug-seeking behavior is more limited than previously appreciated. The use of pharmacological antagonists has indicated that CP-AMPARs underlie the motivational component of cue-drug associative memories formed during drug use [6,7,35], and other work has shown that these receptors contribute to the retrieval of such traces, lowering their activation threshold but not modulating their retention [10]. The molecular mechanisms governing CP-AMPAR incorporation, given the time specificity of their gradual rise only after forced drug discontinuation, are still unknown, although it has been proposed that they might be GluA2-lacking AMPARs and upregulated by NMDAR stimulation and ERK activation [7,36,37]. Continued investigation into the regulation and physiological significance of CP-AMPARs, together with the findings presented here, holds great promise for informing new therapeutic strategies treating the central challenges of individuals who have or are facing substance use disorders.

## ACKNOWLEDGMENTS

This work was supported by a Summer Undergraduate Research Award (SURA) from the Office of Undergraduate Research at the University of Pittsburgh and grants from the National Institutes of Health (MH131717 to OMS).

## AUTHOR CONTRIBUTIONS

OMS led the conceptualization of the study. HA performed the experiments and data analyses, including figure preparation. XH provided electrophysiological training to HA and performed slice preparation for the long-term withdrawal experiments. HA and OMS interpreted the data and jointly wrote the first draft of the manuscript. All authors approved of the manuscript.

